# Valproic acid attenuates hyperglycemia-induced complement and coagulation cascade gene expression

**DOI:** 10.1101/253591

**Authors:** Marina Barreto Felisbino, Mark Ziemann, Ishant Khurana, Camila Borges Martins de Oliveira, Maria Luiza S. Mello, Assam El-Osta

## Abstract

Atherothrombosis remains the leading cause of morbidity and mortality in patients diagnosed with diabetes mellitus, but the molecular mechanisms underpinning this remain unresolved. As the liver plays a major role in metabolic homeostasis and secretion of clotting factors and inflammatory innate immune proteins, there is an interest in understanding the mechanisms of hepatic cell activation under hyperglycemia and whether this can be attenuated pharmacologically. We have previously shown that hyperglycemia stimulates major changes in chromatin organisation and metabolism in hepatocytes, and that the histone deacetylase inhibitor valproic acid (VPA; IUPAC: 2-propylpentanoic acid) is able to reverse some of these metabolic changes. In this study, we used deep transcriptome sequencing to show that VPA attenuates hyperglycemia-induced activation of complement and coagulation cascade genes. These findings reveal a novel mechanism of VPA protection against hyperglycemia, which might improve the therapeutic approaches for diabetes.

### Table of Contents entry (approximately 300 characters excl. spaces) 404 chars

Hyperglycemia induces an inflammatory, prothrombotic state in diabetic patients but the underlying mechanisms remain poorly understood. Felisbino et al, use transcriptomics to identify molecular pathways upregulated by hyperglycemia in hepatocytes, and show that valproic acid attenuates the activation of coagulation and complement genes, which may be useful to reduce cardiovascular risk in diabetes patients.

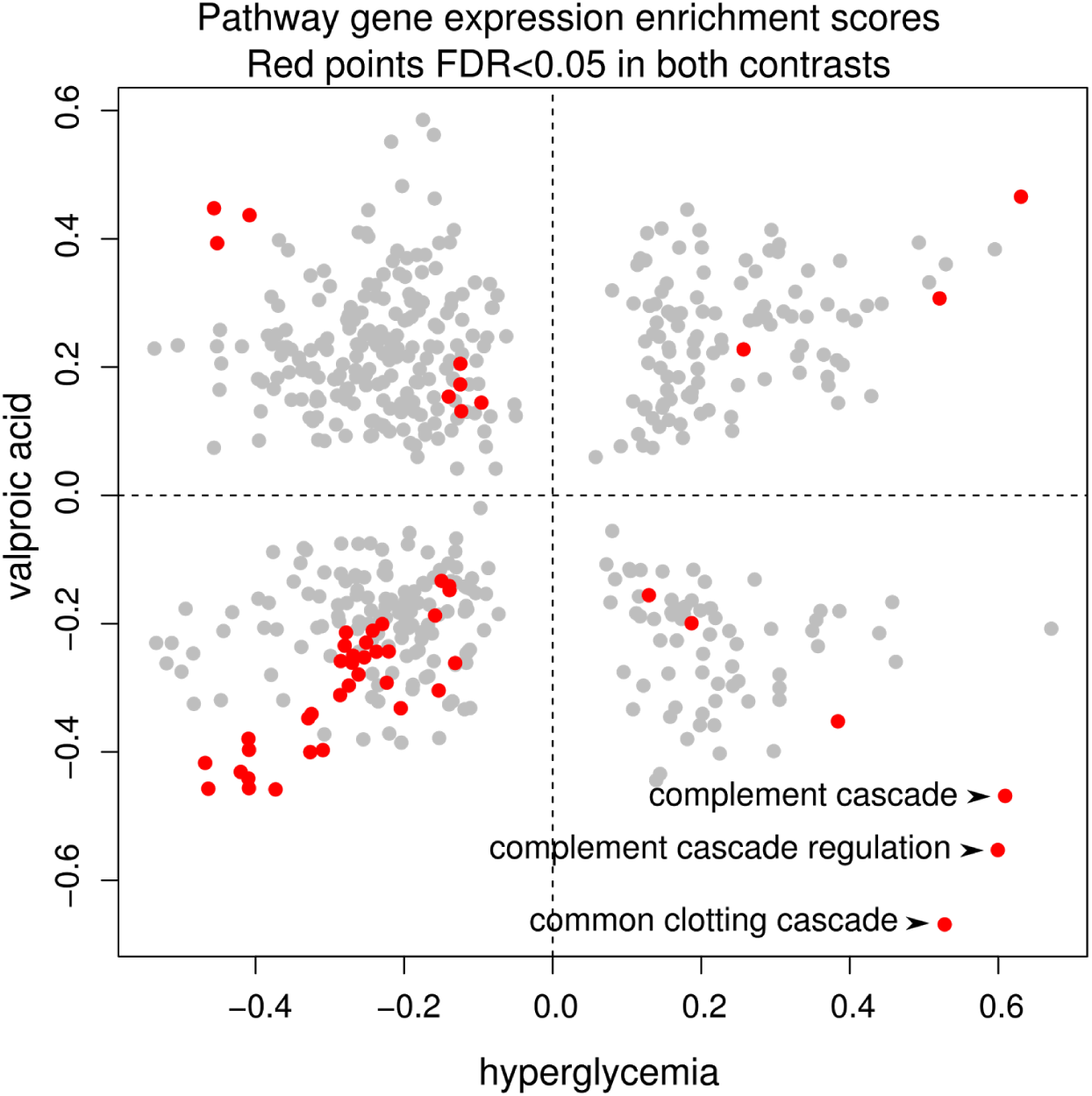

## INTRODUCTION

Diabetes is a multifactorial disorder with several pathways implicated in the development of diabetic micro‐ and macro-vascular complications.**^[1]^** Macrovascular complications include atherosclerosis, cerebrovascular disease and peripheral vascular disease; for which diabetic patients have a two‐ to four-fold greater risk than non-diabetic individuals.**^[2]^** In diabetes, a combination of hyperglycemia, inflammation, oxidative stress and insulin resistance converge to produce a prothrombotic milieu, characterised by endothelial dysfunction, coagulative activation and platelet hyperreactivity.**^[3]^** Several important coagulation pathway proteins are elevated by hyperglycemia *in vivo* including fibrinogen, prothrombin 1 and 2, tissue factor, thrombin-antithrombin complexes, plasminogen activator inhibitor-1, tissue plasminogen activator and complement C3 which contribute to this hypercoagulative state.**^[4,5]^** This prothrombotic state state appears to occur simultaneously with and may be dependent upon chronic low-grade inflammation and oxidative stress observed in diabetes.**^[6]^** Complement cascade proteins are a proposed source of inflammation in diabetes that are produced by monocytes, macrophages and hepatocytes. These proteins function in innate antimicrobial defense primarily through membrane attack and phagocyte recruitment, that are elevated in diabetes and suspected to contribute to diabetic complications.**^[7]^** The liver is the main site of production of circulating coagulation and complement proteins, and responsible for production of bile, cholesterol, decomposition of red blood cells and detoxification of xenobiotics and secondary metabolites. Of key importance for diabetes, liver hepatocytes play a major role in energy homeostasis by storing carbohydrates as glycogen during hyperglycemia and releasing sugars during hypoglycemia. Liver dysfunction, classified as elevated hepatic glucose production during hyperglycemia, is common in type-2 diabetes, and inhibiting this is a major mechanism of action of the widely prescribed glucose lowering drug metformin.**^[8]^** Thus, understanding the molecular mechanisms underpinning the hyperglycemic activation of metabolism, coagulation,complement and other inflammatory pathways in hepatocytes could identify new therapies to reduce the burden of diabetic complications.

At the interface between genetic and environmental factors, epigenetic mechanisms are proposed to play a major role in the development of metabolic disease including diabetic complications.**^[9,10]^** Previous reports have demonstrated that chromatin remodeling and histone acetylation are important mechanisms in diabetes development.**^[11,12]^** The epigenetic component of metabolic/inflammatory disorders has come recently to attention, revealing epigenetic drugs as potential immunomodulatory agents. The recent discovery that histone deacetylase (HDAC) inhibitors (HDACi) have the ability to reduce the severity of inflammatory and autoimmune diseases, including diabetes, in several animal models, has positioned them as alternative anti-inflammatory agents.**^[13]^** Their paradigmatic mode of action has been defined as increased histone acetylation of target genes, leading to higher gene expression; however, recent studies have shown a more diverse mechanism of gene regulation.**^[14-17]^**

Valproic acid (VPA), the most clinically prescribed HDACi, is a fatty acid with anticonvulsant properties used for the treatment of epilepsy and seizures.**^[18]^** Recently, it has been investigated in different disease states as part of a strategy to repurpose clinically approved drugs.**^[19-21]^** There are reports of VPA reducing the blood glucose level and fat deposition in adipose tissue and liver,**^[22,23]^** as well as controlling the insulin production**^[24]^** and gluconeogenesis signaling.**^[23,25,26]^.** It also seems to improve the microvascular complications of diabetes.**^[27,28]^**

We have previously shown that treatment of HepG2 human hepatocytes with the HDACis Trichostatin A (TSA) and VPA attenuated hepatic glucose production, although no significant difference was detected in global chromatin structure and epigenetic landscape. Chromatin alterations promoted by HDACi under hyperglycemia may be a function of the differently regulated nuclear domains and genes rather than of global remodeling.**^[12]^**

Therefore, identification of genes impacted by HDAC inhibition is paramount to a comprehensive understanding of its mechanisms of action and therapeutic target in amelioration of hyperglycemic state.**^[15]^**

We hypothesise that hepatocytes undergo major gene expression alterations when exposed to a hyperglycemic environment as the liver is an organ of critical importance to carbohydrate metabolism. Furthermore, we hypothesised that VPA could attenuate some of the deleterious pathways promoted by hyperglycemia.

In this study, HepG2 cells exposed to high-glucose (HG) were stimulated with VPA. We performed high throughput RNA-sequencing (RNA-seq) to understand transcriptome-wide analysis of of genes and pathways in response to hyperglycemia and VPA. This work identified that complement and coagulation pathways activated by hyperglycemia were strongly attenuated by HDAC inhibition. This work suggests a new avenue of action of VPA that might be relevant to the management of diabetes.

## RESULTS

### Hyperglycemia severely impacts hepatocyte gene expression

In order to understand the effect of high glucose on whole genome hepatic gene expression and function, RNA-seq was performed in HepG2 cells stimulated by hyperglycemic conditions in triplicate. After read alignment and gene expression quantification, statistical analysis of genes and pathways was undertaken. Multidimensional scaling analysis measures the similarity of the samples and projects this on two-dimensions. We observe low glucose (LG) and high glucose (HG) samples clustered into distinct groups (Figure 1A). Statistical analysis showed that HG treatment had a strong effect on HepG2 cells, with 4,259 genes showing differential expression (FDR≤0.05; Figure 1B – red points). This effect is much higher than that reported for high glucose treated monocytes **^[29]^** and endothelial cells **^[30]^** suggesting that hepatic cells are sensitive to changes in carbohydrate supply.

**Figure 1.**
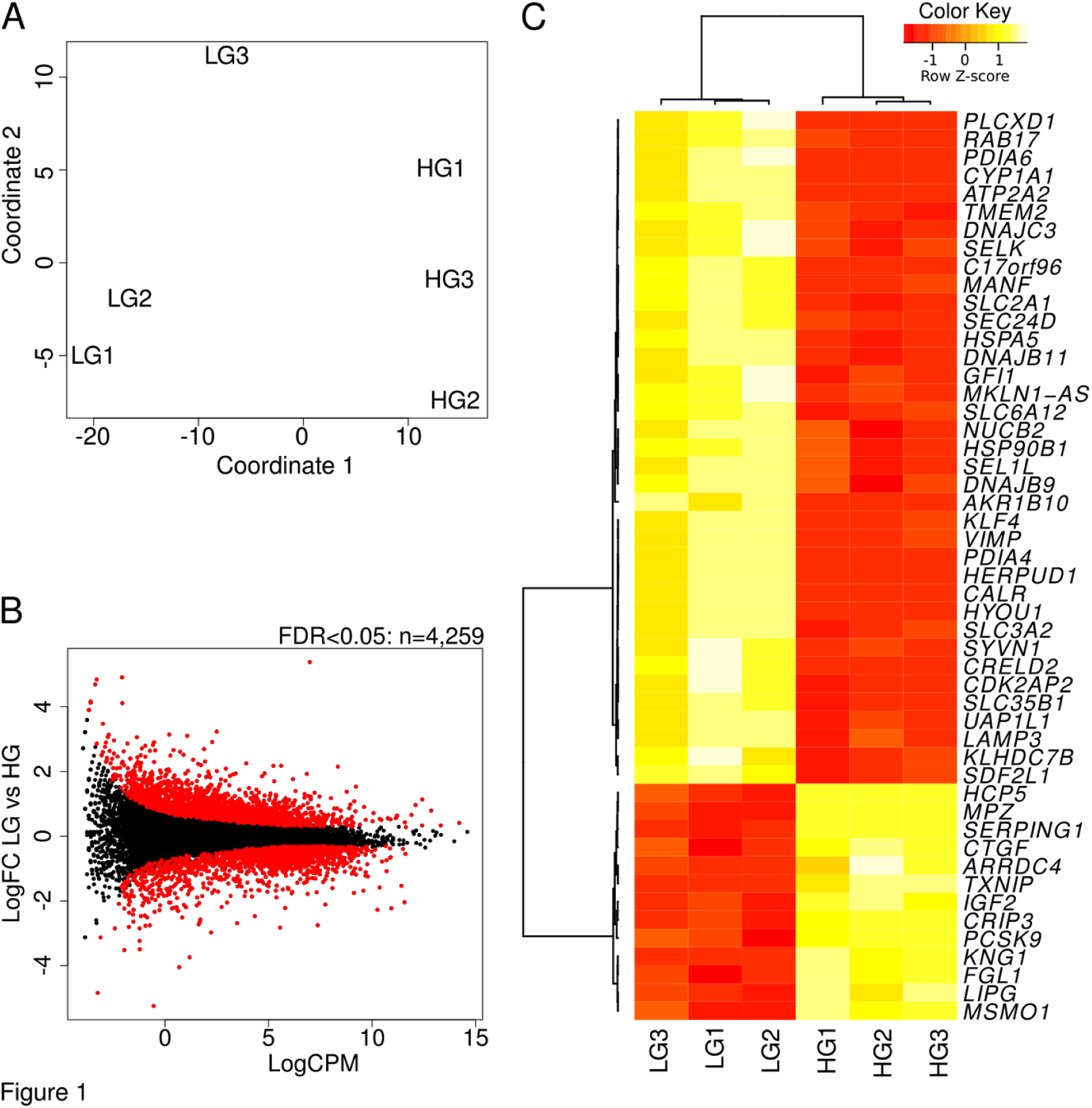
HepG2 gene expression changes in response to high glucose by RNA-seq analysis. (A) Multidimensional scaling analysis of library size normalized counts shows sample clusters based upon treatment on dimension 1. (B) Smear plot depicts the log2 fold change and average expression level log2 counts per million for each detected gene. Genes with differential expression (FDR≤0.05) are highlighted in red. (C) Heatmap of the 50 most significant differentially expressed genes. LG, low glucose ‐ normoglycemic condition. HG, high glucose – hyperglycemic condition.

The top 50 differentially expressed genes by significance are shown in heatmap form (Figure 1C). Some of the up-regulated genes are: *TXNIP*, a known hyperglycemia inducible gene that is highly abundant in HepG2 cells (TXNIP protein inhibits the normal function of thioredoxin leading to accumulation of ROS); *SERPING1*, which encodes C1 esterase inhibitor, and is involved in inhibition of complement cascade; *MPZ*, that encodes a structural component of the myelin sheath and is thought to be specific to nervous system tissues; *MSMO1*, that encodes a protein involved in cholesterol biosynthesis; *PCSK9*, whose protein acts in binding to and degrading low-density lipid receptors; *IGF2,* that encodes an insulin-like growth factor, the central regulator of somatic growth and cell proliferation. Some of the down-regulated genes are: *MANF*, whose protein promotes survival of dopaminergic neurons, possibly playing a role in ER stress response; *HSPA5* (aka *GRP78*), which encodes a glucose sensing protein, whose expression is upregulated by glucose starvation and is also present in the ER; *CALR*, that encodes calreticulin, a major calcium storage protein in the ER; *DNAJB11*, which encodes yet another ER localized protein that acts as a chaperone for a number of partners. The glucose transporter gene *SLC2A1*, encoding GLUT1 was also strongly down-regulated.

Gene Set Enrichment Analysis (GSEA) was used in order to understand pathways regulated by hyperglycemia. From 575 REACTOME gene sets considered, 34 were upregulated and 139 were down-regulated (FDR≤0.05). The top 20 gene sets by significance in the up and down-regulated directions are shown (Figure 2A). Down-regulated gene sets included those associated with extracellular matrix interactions, chaperone function, calnexin/calreticulin cycle, N-glycan trimming and peptide chain elongation (Figure 2A), while gene sets upregulated in response to hyperglycemia included cholesterol biosynthesis, complement cascade and fibrin clotting cascade (Figure 2B, C, D). These findings show a distinctive response of hepatocytes to hyperglycemia.

**Figure 2.**
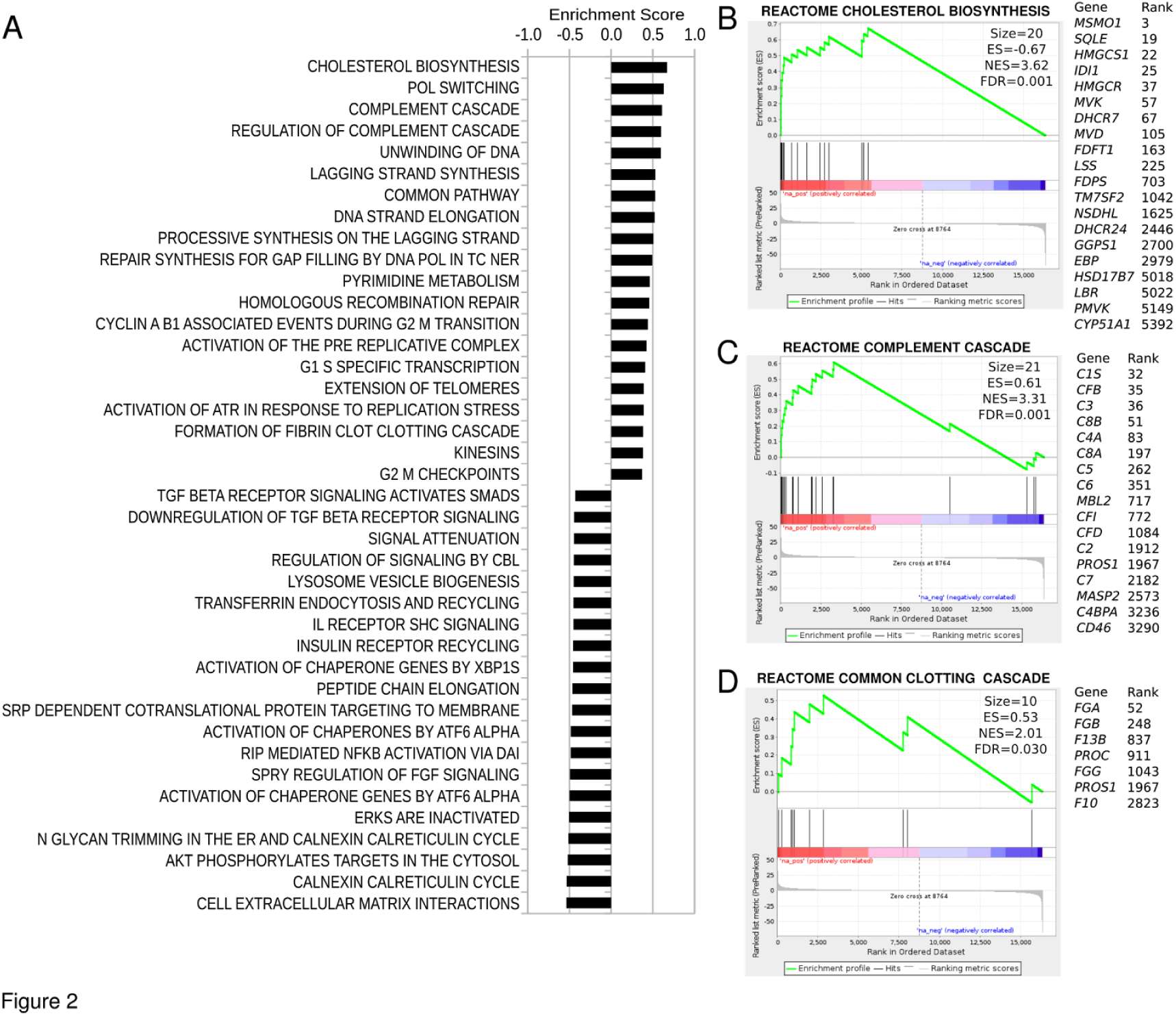
Gene sets differentially expressed in response to high glucose. (A) Top 20 REACTOME pathways with differential expression selected by statistical significance, determined by pre ranked gene set enrichment analysis (GSEA-P). (B, C, D) Enrichment plots show upregulation of cholesterol biosynthesis, complement cascade and common clotting cascade in response to hyperglycemia. All pathways GSEA FDR <0.05.

### VPA treatment attenuates the expression of hyperglycemic response genes

Given that hyperglycemia induces major changes to the hepatocyte transcriptome and activates pathways relevant to cardiovascular health (such as cholesterol metabolism and complement/clotting cascades) and our previous work shows that VPA attenuates hepatic function, we hypothesised that VPA might inhibit hyperglycemic gene expression signatures.

Multidimensional scaling analysis shows that samples cluster based on treatment group. Untreated samples (LG, HG) are clearly separated from VPA-treated samples (LGV, HGV); and normoglycemic samples (LG, LGV) are separated from hyperglycemic ones (HG, HGV) (Figure 3A). Smear plot shows that 7,802 genes were altered in expression due to VPA treatment under hyperglycemia (Figure 3B). This plot also shows genes with initially low expression were upregulated after VPA treatment; on the other hand, genes initially highly expressed were down-regulated. Heatmap of top 50 genes by significance shows that the majority were upregulated (Figure 3C).

**Figure 3.**
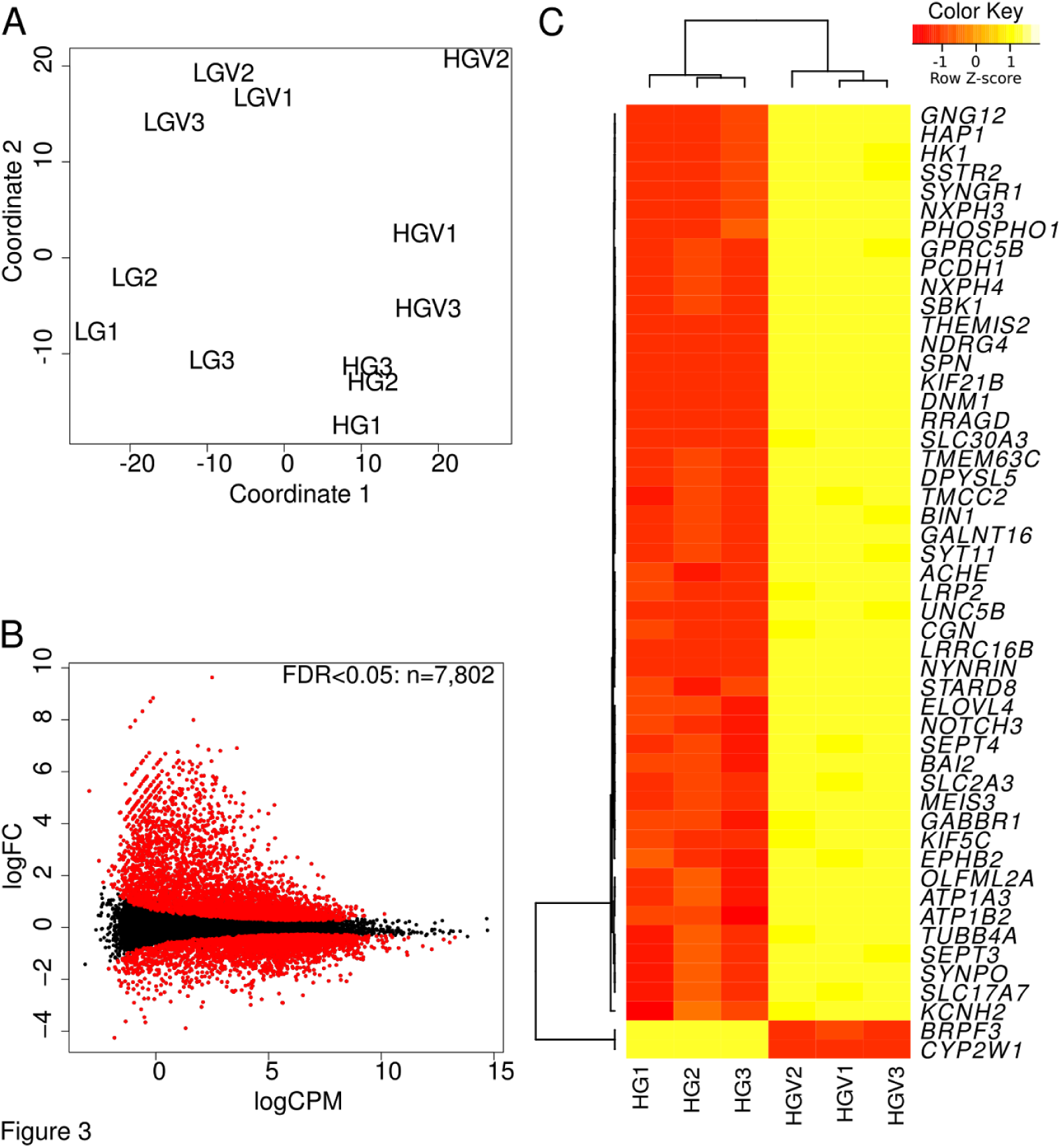
Hyperglycemic HepG2 gene expression in response to VPA. (A) Multidimensional scaling analysis of library size normalized counts shows samples cluster based upon treatment groups. (B) Smear plot showing the effect of VPA treatment on hyperglycemic HepG2 cells. Genes with differential expression (FDR≤0.05) are highlighted in red. (C) Heatmap of the 50 most significant differentially expressed genes responding to VPA. LG, low glucose – normoglycemic condition. LGV, normoglycemic condition followed by VPA treatment. HG, high glucose – hyperglycemic condition. HGV, hyperglycemic condition followed by VPA treatment

The top 20 gene sets by significance in the up‐ and down-regulated directions are shown (Figure 4A). Gene sets upregulated included those related to function of neurons including potassium channels, neurotransmitter receptor, L1-type/ankyrins interactions. Down-regulated gene sets included common pathway of fibrin clot formation, complement cascade and genes involved in protein synthesis. Clotting and complement cascade genes were down-regulated by VPA in hyperglycemic condition (Figure 4B,C). The regulation of all genes in response to glucose and VPA was visualised on a two dimensional rank-rank plot (Figure 4D). We observe that overall, genes are distributed relatively evenly among the four quadrants.

**Figure 4.**
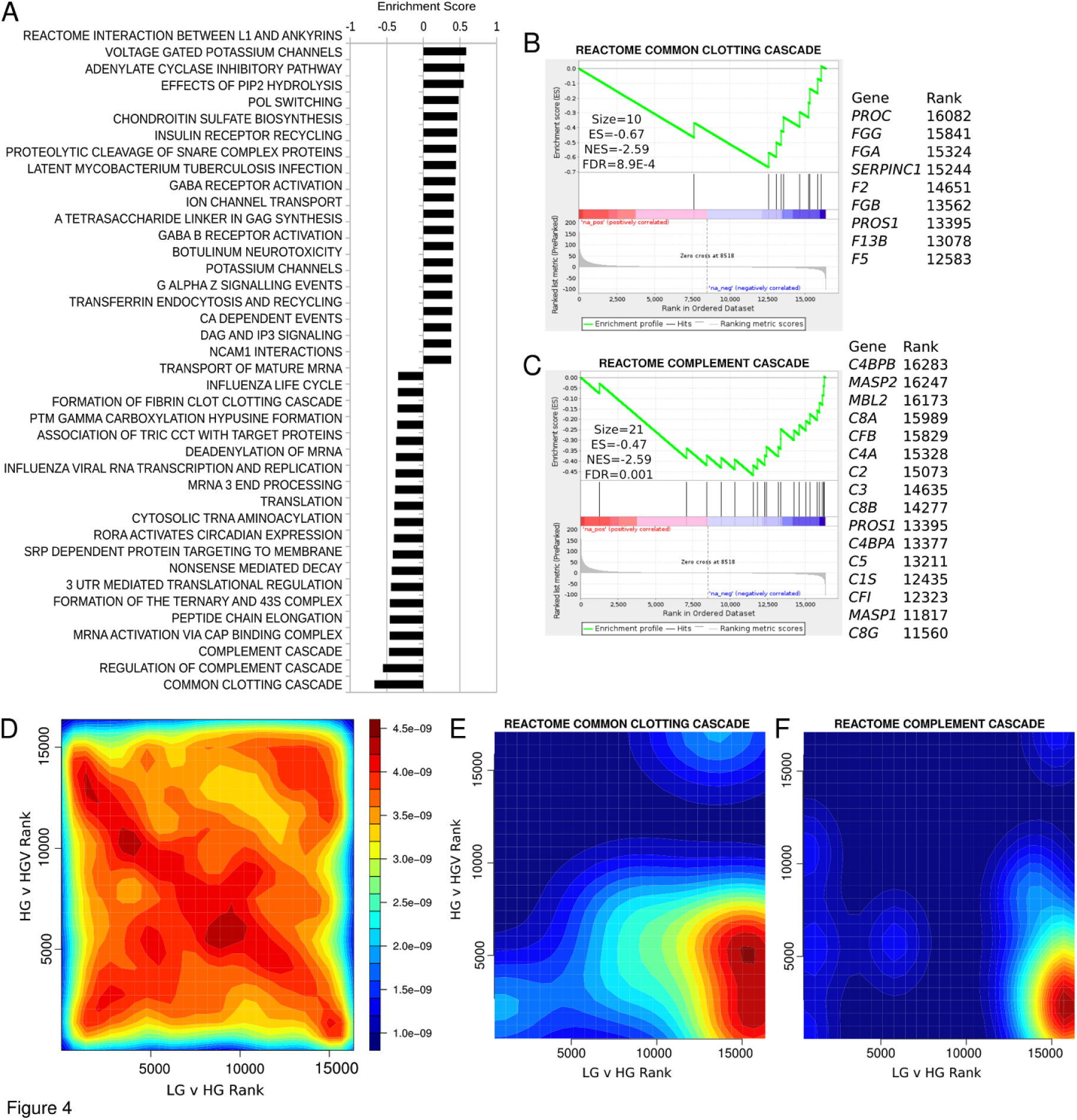
VPA attenuates expression of clotting and complement pathways in hyperglycemia-treated HepG2 cells (A) Top 20 REACTOME pathways with differential expression in response to VPA under hyperglycemia determined with preranked gene set enrichment analysis (GSEA-P, FDR <0.05). (B,C) Enrichment plot shows downregulation of common clotting and complement cascades in response to VPA. (D) Rank-rank plots of gene expression changes for all detected genes and (E,F) specifically for genes in the common clotting and complement cascade pathways.

Using rank-rank visualisation of clotting and complement cascade genes we observed coordinated upregulation of these genes with hyperglycemia and attenuation by VPA (Figure 4E,F). The FDR corrected MANOVA p-values for the two-dimensional association were 2.0E-4 and 1.5E-7 for clotting and complement cascades respectively.

To validate some differentially expressed genes from the RNA-seq findings, we cultivated HepG2 cells under hyperglycemic conditions prior to treatment with VPA as above followed by quantitative reverse transcription PCR (RT-qPCR). The selected genes included those involved in complement (*MASP2*) and clotting cascade (*FGA*, *PROC*, *FXIIB*). Selected genes upregulated by hyperglycemia according to the RNA-seq results, and confirmed by real-time PCR, were attenuated by VPA stimulation in agreement with RNA-seq (Figure 5). Taken together our results show that VPA attenuates hyperglycemia-induced expression of complement and clotting genes.

**Figure 5.**
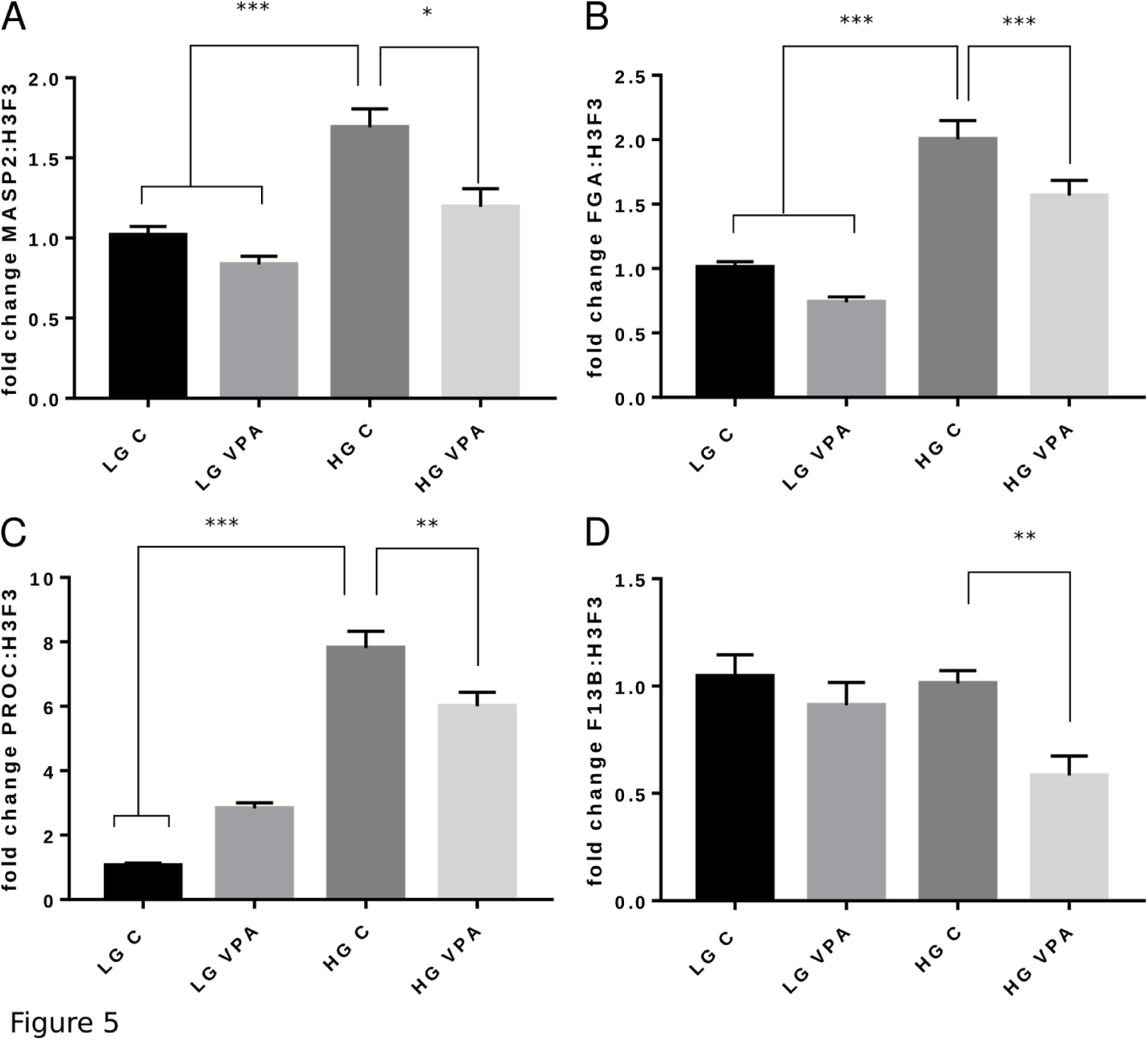
VPA stimulation attenuates the expression of complement (*MASP2*) and clotting (*FGA*, *PROC, F13B*) pathways genes as validated by qPCR. LG, low glucose – normoglycemic condition. LGV, normoglycemic condition followed by VPA treatment. HG, high glucose – hyperglycemic condition. HGV, hyperglycemic condition followed by VPA treatment. *, p <0.05; **, p <0.01; ***p <0.001 as depicted by one-way anova analysis.

## DISCUSSION

Metabolic syndromes and associated cardiovascular complications are a major health burden, but still, we have limited information on how deleterious stimuli such as hyperglycemia impact gene regulation in different organs, tissues and cells of the body. As the liver plays a major role in energy homeostasis, we hypothesised that hepatocytes would demonstrate robust gene expression changes in response to hyperglycemia, some of which could be deleterious to cardiovascular health. Furthermore, we hypothesised that HDAC inhibition via VPA could reverse or attenuate some of these gene pathways.

We used high throughput RNA sequencing (RNA-seq) for its unbiased ability to detect all expressed genes with greater sensitivity and accuracy than gene expression microarrays. With appropriate bioinformatics tools, regulatory events to genes and pathways (sets of genes) can be pinpointed in a way that is more efficient than single-gene assays. These tools were applied to identify genes and pathways that respond to hyperglycemia and/or VPA in hepatocytes.

The major gene sets upregulated by hyperglycemia were related to cholesterol metabolism, DNA replication and complement cascade and clotting cascades. The observation of elevated expression of clotting and complement factors is consistent with reports of these proteins being elevated in patients with diabetes. Interestingly, of these pathways, only complement cascade and clotting cascades were attenuated by VPA, as demonstrated by our two dimensional pathway analysis.

The complement system is a central component of innate immunity against microorganisms and modulator of inflammatory processes; it comprises a complex and tightly regulated group of proteins involving various soluble and surface-bound components. Depending on the activation trigger, the complement cascade follows one of three pathways: classical, lectin or alternative.**^[31]^** Although these pathways differ in their mechanisms of target recognition, all converge in the activation of the central component C3. This process is followed by C5 cleavage and the assembly of the pore-like membrane attack complex, MAC. Important chemoattractants and inflammatory mediators are produced by the enzymatic cleavage of C3 and C5, which leads to the release of anaphylatoxins C3a and C5a.**^[32]^**

Together with complement, the coagulation system is the major blood-borne proteolytic cascade.**^[33]^** Upon activation of the coagulation cascade, a sequential series of serine protease-mediated cleavage events occur. Thrombin is activated from its zymogen prothrombin and then catalyze the polymerization of fibrin by cleaving small peptides from its subunits. This way, soluble fibrinogen is converted into insoluble fibrin, which allows the clot formation.**^[34]^** Thrombin also plays a key role in amplifying the cascade by feedback activation of coagulation factors.**^[35]^** Other components as circulating red and white blood cells, and platelets are incorporated into the clot structure. In addition, factor XIIIa, which is also activated by thrombin, provides further structural stability by cross-linking with fibrin.**^[36]^** In this context, weak clots are more susceptible to fibrinolysis and bleeding, while resistant clots may promote thrombosis.**^[34]^**

Coagulation and complement cascades share a common evolutionary origin **^[33]^** and their interplay is highlighted by C3, C4, C5a and FB complement protein presence in thrombus,**^[37]^** Other components as circulating red and white blood cells, and platelets are also procoagulation enzymes thrombin and IXa, Xa, XI factors might activate complement cascade.**^[38]^** Moreover, MASP2, a component of lectin complement activation, is capable of cleaving coagulation factors prothrombin in thrombin, fibrinogen, factor XIII and thrombin-activatable fibrinolysis inhibitor *in vitro*. **^[39,40]^** Thus, understanding the crosstalk between these pathways has fundamental clinical implications in the context of diseases with an inflammatory and thrombotic pathogenesis, in which complement–coagulation interactions contribute to the development of complications.**^[41]^**

Liver, mainly hepatocytes, is responsible for the biosynthesis and secretion of the majority of complement and coagulation components. Furthermore, the promoter regions of these components are controlled by several common liver-specific transcription factors like HNFs and C/EBP.**^[42]^** Thomas and co-workers **^[43]^** compared the genome-wide binding of Fxr and Hnf4α in mouse liver and characterized their cooperative activity on binding to and activating target gene transcription. Genes bound by Fxr and Hnf4α are enriched in complement and coagulation cascades, as well as in pathways related to drug metabolism. Furthermore, these transcription factors are involved in gluconeogenesis and glycogenolysis gene expression.**^[44,45]^** Thus, a common transcription factor network may be controlling the immune system and metabolism pathways.

The participation of complement in metabolism and metabolic disorders has recently received increasing scientific attention. Earlier studies demonstrate higher plasma C3 levels in diabetic patients compared to healthy individuals.**^[46,47]^** Complement activation has recently been reported to be involved in arterial hypertension, and may explain the similar clinical features of malignant nephrosclerosis and atypical hemolytic uremic syndrome.**^[48]^** Increased complement gene expression has also been associated with adipocyte insulin resistance, waist circumference, syndrome **^[35]^** and triglyceride levels.**^[49,50]^** MASP-1 and MASP-2 levels are found at significantly higher levels in children and adults with T1DM than in their respective control groups, whereas these proteins levels decreased when glycemic control improved.**^[51^]**

Metabolic syndrome, including diabetes mellitus, is also frequently associated with a procoagulant state, in which the clotting system is switched toward a prothrombotic state, involving reduced fibrinolysis, increased plasmatic coagulation, and platelet hyperactivity.**^[35,52,53]^** Intensive glycemic control with insulin reduces the impact of this procoagulant state by affecting components of the clotting pathway.**^[53]^** Abnormalities in the coagulation and fibrinolytic systems contribute to the development of cardiovascular complications in patients with metabolic and consistent lowering of clotting factors are used for the treatment of acute cardiovascular syndromes.**^[54]^**

As a prime initiator and important modulator of immunological and inflammatory processes, the complement system has emerged as an attractive target for early and upstream pharmacological intervention in inflammatory diseases.**^[55]^** In this context, repurposing clinically approved drugs such as VPA provides a time‐ and cost-effective alternative.

## CONCLUSIONS

In this work we could, for the first time, associate HDAC inhibition with complement and coagulation gene expression modulation, confirming that coagulation and complement cascade genes were upregulated by hyperglycemia and that these can be attenuated with VPA. If confirmed by pre-clinical and clinical *in vivo* assays, these findings have the potential to improve the therapeutic approaches for diabetes treatment.

## EXPERIMENTAL SECTION

### Cell culture

HepG2 cells from ATCC at passage 9 were maintained in Dulbecco’s modified Eagle’s medium (DMEM) basal glucose (5.5 mM) (Gibco, Carlsbad, USA) supplemented with 10% fetal bovine serum (GE Healthcare, Chicago, USA) and penicillin and streptomycin (Gibco) (working dilution: 100 IU and 100 μg/mL, respectively). The cells were cultivated for 48 h in normoglycemic (LG) or hyperglycemic (HG) medium, containing D-glucose (Sigma, St. Louis, USA) up to a final concentration of 20 mM. Cells under LG and HG conditions were then treated with 1.0 mM VPA (Sigma) for another 12 h and compared with the respective untreated controls.

### RNA isolation

Cells were disrupted with TRIzol (Qiagen, Hilde, Germany). RNA was isolated from TRIzol homogenates using the Direct-zol Kit (Zymo Research, Irvine, USA). RNA was quantified on Qubit (Thermo Fisher, Waltham, USA) and its quality was evaluated on the MultiNA bioanalyzer (Shimadzu, Kyoto, Japan).

### mRNA sequencing

NEBNext^®^ Poly(A) mRNA Magnetic Isolation Module was used to enrich mRNA from μg of total RNA. We used the NEBNext^®^ Ultra^TM^ Directional RNA Library Prep Kit for Illumina^®^ (San Diego, USA) to generate barcoded libraries. Libraries were quantified on the MultiNA bioanalyzer (Shimadzu) and pooled to equimolar ratios for sequencing.

Cluster generation was performed at a concentration of 10 pM (TruSeq SR Cluster Kit v3-cBot-HS) and the flow cell was run on Illumina HiSeq2500 generating 50 nt reads at AGRF Melbourne. Sequence data has been deposited to NCBI Gene Expression Omnibus under accession number GSE109140.

### Bioinformatics analysis

#### mRNA-seq data processing

Low quality bases (Qscore < 20) were removed from the 3’ end with FASTX Toolkit v0.0.13. Trimmed reads less than 20 nt were also discarded. The human genome sequence and annotation (GRCh37.75) set were downloaded from the Ensembl website (http://www.ensembl.org/info/data/ftp). Reads were aligned using STAR version 2.3.1p_r359.**^[56]^** Sorted bam files were generated with SamTools (version 0.1.19-44428cd).**^[57]^** A gene expression count matrix was generated with featureCounts v1.4.2 **^[58]^** using a map quality threshold of 10. Genes with an average of fewer than 10 reads per sample were omitted from downstream analysis.

#### Statistical analysis of gene expression

EdgeR version 3.6.8 and limma version 3.20.9 were used to perform statistical analysis.**^[59]^** False discovery rate controlled p-values (FDR)≤0.05 were considered significant.

#### Pathway analysis

Gene expression of pathways was analyzed with GSEA-P using the classic mode.**^[60]^** A differential abundance score was obtained for each gene by dividing the sign of the fold change by the log10(p-value). This score was used to rank genes from most up-regulated to most down-regulated as described previously.**^[15]^** Curated gene sets were downloaded from MSigDB.**^[61]^**

#### Integrative analysis

To understand the correlation between effects of VPA and hyperglycemia on global gene expression, we generated a rank-rank density plot of each detected gene. Genes were ranked as above and plotted using the filled contour plot function in R. Significance of two dimensional enrichment of gene sets away from a uniform distribution was calculated Manova test of ranks in R as described previously.**^[62]^**

### Real time quantitative PCR

To validate some differentially expressed genes from the RNA-seq findings, we repeated the experiment using the same conditions of cell culture and treatment, isolated total RNA using the RNeasyMini Kit (Qiagen, Hilden, Germany) and prepared cDNA using the High-Capacity cDNA Reverse Transcription Kit (Applied Biosystems, Waltham, MA). Real-time PCR was performed using an Applied Biosystems 7500 Real Time PCR system following standard protocols and TaqManGene Expression assays (Applied Biosystems) for complement (*MASP2* (Hs00373722_m1)) and coagulation cascade (*PROC* (Hs00165584_m1), *FGA* (Hs00241027_m1), *F13B* (Hs01055646_m1)) genes.

Target gene expression was normalized to the expression of H3F3 (Hs02598544_g1) Relative quantification was achieved with the comparative 2-ΔΔCt method as described previously.**^[63]^**

## ACKNOWLEDGEMENTS

We acknowledge the use of Illumina sequencing at AGRF (and the support it receives from the Commonwealth of Australia). This work was supported by the São Paulo Research Foundation (FAPESP, grants no. 2010/50015-6, 2012/03238-5, 2014/10198-5, and 2015/10356-2) and the Brazilian National Council for Research and Development (CNPq, grant no. 304668/2014-1).

